# A low-cost protocol for reconditioning of deep-brain neural microelectrodes with material failure for electrophysiology recording

**DOI:** 10.1101/2024.01.17.576102

**Authors:** Leila Rezayat, Mohammad Hossein Ghajar, Alireza Naji, Jalaledin Nourozi, MohammadReza Abolghasemi Dehaqani, Ehsan Rezayat

## Abstract

Up to now, a large variety of neural microelectrodes are developed. Although there are a lot of published works in this area, the majority of them are about the fabrication methods and rarely discuss the reconditioning procedure or how to re-use the electrodes that their performance is decreased because of material failure. In this paper, firstly, it is answered that why the performance of neural microelectrodes decreases. Secondly, a general low-cost protocol for reconditioning and re-using electrodes are proposed that can be utilized for the most types of electrodes with material failure. Lastly, the proposed reconditioning protocol is applied experimentally to single-site tungsten microelectrodes to demonstrate the effectiveness of the protocol. Neural signal recording Results clearly indicate that a large number of electrodes can be reconditioned well.

## Introduction

During the last decades, deep brain neural recording electrode technologies have been developed rapidly, starting from microwire electrodes, and then, other types such as Michigan and Utah array, and most recently, flexible electrodes (Hong & Lieber, 2019). However, they are fabricated well, still suffer from failure and their performance will decrease after several usages. Barrese et al. defined failure simply as the absence of extractable action potentials for all electrodes (channels) on an array(Barrese et al., 2016), and then generally established four discrete failure mode categories: biological, material, mechanical, and unknown.

Biological failures are those related to the foreign body response of the tissue to the sensor including bleeding, cell death, hardware infection, meningitis, gliosis, or meningeal encapsulation and extrusion. Material failures are related to inherent design flaws or material degradation, for example, broken electrode tips, or insulating materials leakage, cracks, and delamination. Mechanical failures are related to physical factors that move the sensor from its desired location or damage the hardware enough to prevent recordings such as bundle damage, connector damage, and mechanical removal. Unknown failures are signal loss not directly attributable to one of the three mentioned mechanisms(Barrese et al., 2016).

Several researchers have studied the material failure mechanism of neural microelectrodes (Barrese et al., 2016; Gilgunn et al., 2013; Prasad et al., 2012, 2014; Schmidt et al., 1988; Simeral et al., 2011). Their results indicate that insulating material failure is the most significant failure mode among all due to the graduate decrease in impedance and signal quality. Particularly, material failure modes in this research include corrosion, cracking, bending of recording sites, and delamination or cracking of the insulating polymer materials(Kozai et al., 2015).

In spite of all these publications and studies, there are only few non-significant researches on how to fix a material failure or reconditioning electrodes to be able to utilize them efficiently for a longer time. For example, Li et al. described a method for the construction, repair, and recycling of tungsten-in-glass micro-electrodes (Li et al., 1995). Their method includes either removing excess glass insulation from the microelectrode tip or fine adjustment and reshaping of the exposed electrode tip. Their approach enables the controlled shaping of the electrode tip during preparation and/or reshaping the electrode tip or glass insulation of damaged electrodes. Verhagen et al. presented a cost-effective and simple procedure to restore initial high impedance of epoxy-insulated tungsten microelectrodes by dipping the electrodes in epoxy followed by curing (Verhagen et al., 2003).

Here, after a brief introduction and literature review, firstly, different causes of the failure of neural microelectrode are discussed in detail. Particularly, a circuit model is utilized for the insulated electrode to describe better the material failure and impedance decrease. Secondly, a general low-cost reconditioning protocol is proposed to re-use a wide range of different types of electrodes with material failure. Then, to determine how much the proposed reconditioning protocol effectiveness is, a single-site tungsten microelectrode type as a case study is examined. In continue, the electrophysiology recording results for the case study are presented and discussed. Finally, it is concluded in the last section.

## Causes of electrode failure

As it is mentioned before, failure modes are categorized into four groups that are biological, material, mechanical, and unknown (Barrese et al., 2013). Among all these, material failure is the most likely reason for the failure of neural microelectrodes. More specifically, it is the insulating failure that mostly causes a decrease in impedance and signals intensity. An insulating polymer material can fail due to leakage, delamination, and cracking. Another aspect of material failure is related to the physical damages such as breaking, cracking, and bending of electrode tips and recording sites (Kozai et al., 2015).

The quality of a neural signal either a spiking activity or a local field potential (LFP) in an electrophysiological recording depends on the impedance of the electrode. As a result, any change in impedance will affect the recorded neural signal quality. Figure 1A demonstrate The impedance of the conical electrodes strongly depends on the material of the conductor, the diameter of the electrode in the tip and in the cylindrical part, and the intensity of the acute angle of the tip. Robinson has represented an equivalent circuit for an implanted metal electrode next to a neuron cell which is generalizable for most types of neural electrodes(Robinson, 1968). The equivalent circuit is demonstrated in Figure 1B. *C*_*s*_ is all the shunt capacitance to the ground from the tip to the input of the amplifier. This includes the capacitance from the metal of the electrode to the bath through the insulation as well as the accumulated capacitance of all the connectors and (shielded) wires leading from the preparation to the amplifier (Robinson, 1968). If we assume that the connectors and wirings are ideal, then the insulator will be the dominant portion of *C*_*s*_ Open recording site of the electrode (i.e. the part of the electrode with no insulation and directly in contact with electrolyte) is modeled by *C*_*e*_ and *R*_*e*_ representing an electric double layer capacitance with some current leakage.

**Figure 1:**
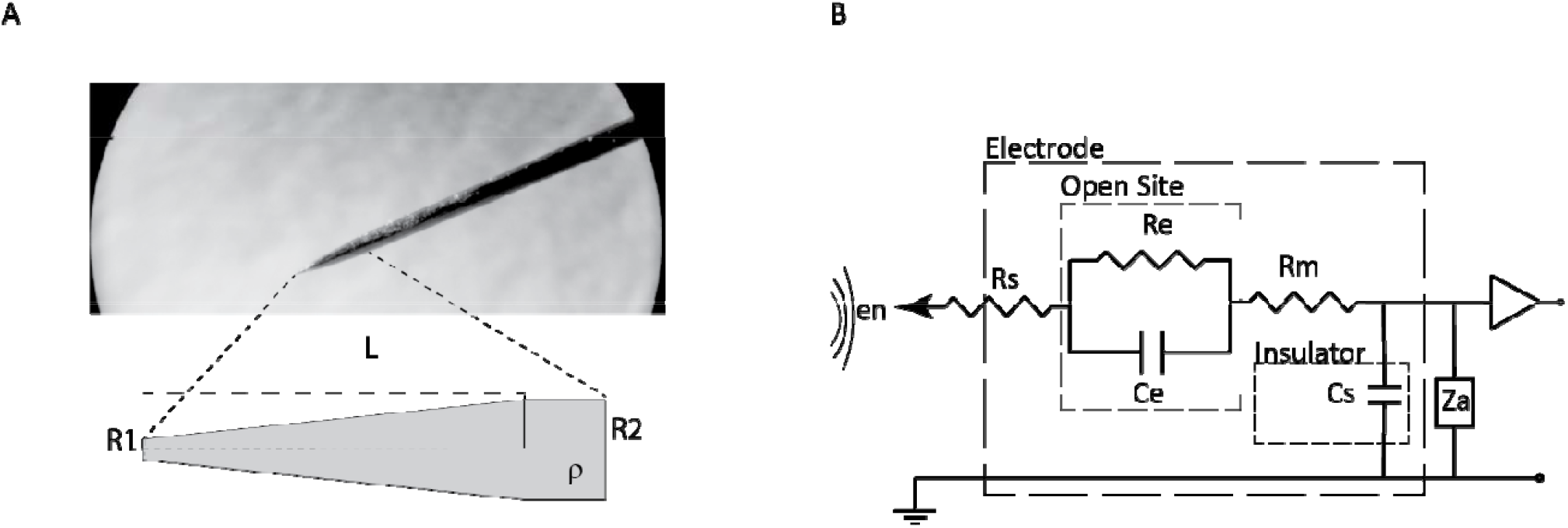
schematic of electrode tip and its equivalent electrical circuit. A) A picture taken with an optical microscope of the tungsten electrode tip and the schematic of the electrode tip. The electrode impedance is determined based on the size of the electrode tip and electrode radius and the distance between these two parts of the electrode, and its value is calculated with ——. : Radius of the electrode tip, : Radius of the electrode cylinder, : Distance between tip and cylinder, : Electrical resistivity and conductivity. B) equivalent electrical circuit of electrode with : Amplifier impedance, : Shunt capacitance, : Metallic resistance of th microelectrode, : Electric double layer capacitance at the interface of the metal tip and the electrolyte solution, : Leakage resistance, : Saline bath resistance (spreading resistance), : Potential created by the extracellular currents.

In the case of recording sites which are covered by an insulating layer, the open site block in Figure 1B should be replaced by the insulator capacitance, or simply considering a relatively infinite amount for *R*_*e*_. In other words, there would be a neglectable leakage current resistance (*R*_*e*_ ≈ 0) and the insulator capacitance is divided into two parts, i. recording area (*C*_*e*_) and ii. Rest of the covered area (*C*_*s*_). It is common in the literature to merge these two parts of insulator capacitance for simplicity while the recording site is covered. That means they consider only *C*_*s*_ which representing the capacitance of the whole insulator including the area of recording sites as well.

By assuming non-open recording sites and neglecting *R*_*m*_ and *R*_*S*_ due to the small amount, the equivalent impedance of the electrode for any specific frequency ω is defined as Equation 1:

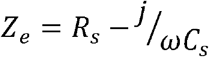

From Equation 1 it can be clearly seen that the *C*_*s*_ is the major parameter in determining the amount of the electrode impedance (*Z*_*e*_). Therefore, any failure in insulator dielectric properties will cause a failure in the electrode impedance and so, in neural signal sensing performance. For example, when delamination occurs, or a cracking happens, or saline molecules diffuse in the electrode insulator, *C*_*s*_ changes significantly, and then, *Z*_*e*_ will be in out of demand range.

On the other hand, physical damages such as breaking, cracking, and bending of electrode tips and recording sites also can change the electrode impedance due to changes in the insulator capacitance properties. Besides, in most cases, a physically damaged electrode will be basically not useable anymore. For instance, a bended tip cannot penetrate into a brain, or a broken tip may cause some brain tissue damages during its movement. Physical damages occur while the electrode is treated improperly during handling or setup. And sometimes it will be damaged when it hits the internal surface of the skull during a deep brain penetrating.

## Reconditioning protocol

Here, a protocol is presented for reconditioning electrodes with material failure. Based on the previous section, we know now that this type of failure is related mainly to the damaged insulator. So, the protocol is aiming to repair the damaged insulator coating. Re-annealing the electrode is the first and most straightforward step for reconditioning it. The re-annealing temperature is required to be a bit above glass temperature to let the insulator polymer chains to be reconfigured and to repair some defects such as delamination and cracks. This method is effective for a large percentage of damaged electrodes and after that, electrodes can be re-used.

If only re-annealing the electrodes does not repair them, then, re-coating of the insulator can fix the out of tune impedance failure. In this method, a new insulation coating will apply over the current layer. However damaged electrode could benefit from this method, it comes with some drawbacks such as diameter increase. If the increase in diameter is not tolerated, then the current existing layer should be removed before the replacement of a new layer. Removing the current layer based on the type of the electrode could be varied. It is possible to be removed by scratching, etching, burning, or etc.

In some cases, a more complicated reconditioning procedure is required in addition to the re-annealing or the re-coating method to fix a damaged electrode. Regarding the type of electrodes, some of them need to be trimmed, some to be re-sharpen, some to be pulled or stretched mechanically. This method can fix some electrodes with physical damages. Our proposed protocol is summarized in Figure 3. In the next section, a regular tungsten tip is reconditioned as a case study to describe the protocol in detail.

## Case study

A single-site neural microelectrode is studied here to demonstrate the effectiveness of our proposed protocol. The impedance of tungsten electrodes will decrease after several uses without any physical failure. To recondition these electrodes, re-annealing them at 160 □for 2 hours is usually sufficient. If the impedance still remains low, the second method of re-coating is applied. Epoxy EL-413 in combination with Hardener HA-23 and Accelerator AX-10 (an Iranian company) is used as a new insulator layer. To prepare the coating solution, first, 0.5 wt% of the hardener is heated to 60 °C. Then, 47.1 wt% of the accelerator is added and mixed gradually. Finally, 52.4 wt% of the epoxy is added and stirred. Another method for insulation is to use polyethylene terephthalate. This material is heated up to the melting temperature of 250 □ to make it liquid and ready to use. After that, the electrode is rinsed with distilled water followed by ethanol and then dipped in the coating solution. Next, the electrode is pulled out slowly and placed in a holder vertically so that its tip is pointing down. Then, a hot air flow, possibly generated by a hot gun, is used to smooth the surface of the coating. Finally, the electrodes are located in an oven for 2 hours at 160□ to be annealed.

If there is a physical fracture in the electrode, such as a bent tungsten tip, it must be treated before the re-annealing or re-coating operation. For the folded tip, chemical etching and trimming were used for both small and large folded tips. After removing the creased defects, the insulating coating was also removed by shaving with a sharp tool such as a cutter. Then, the tip was sharpened again. For this purpose, a 10 M potassium hydroxide (KOH) base was used as the etching solution, and a sinusoidal voltage with an amplitude of 10 V and a frequency of 10 Hz was applied to the cathode/anode, as shown in Figures 2a and 2b. The setup provides both manual and automatic modes for adjusting the position of the electrode in the solution. This position adjustment is done by moving the KOH solution between the two containers. To begin, we place the electrode in the main chamber and front of the camera lens using a clamp so that its tip is visible in the image. Then, we bring the solution level closer to its head using manual control. After that, we activate the automatic control. The automatic control activates the motor for the necessary time in the direction where the height of the solution rises. This time of increasing the solution should be enough for the water level to reach the electrode resin coating, which should be adjusted by the person present in the program. At this moment, the operation of raising the solution stops, and immediately the motor is activated in the direction of lowering the solution until it reaches the starting point. This action of going up and down is repeated continuously. This helps the oxides fall from the metal surface and speeds up the removal process. It is also possible to control the acute angle of the electrode tip, with smaller angles at lower rates of solution surface change and larger angles at higher rates. The pump motor controller circuits are shown in Figure 3. The procedure ends when the resolution of the electrode tip is observed in the camera. The electrode is then thoroughly washed with distilled water and finally covered in the same manner as described above.

**Figure 2:**
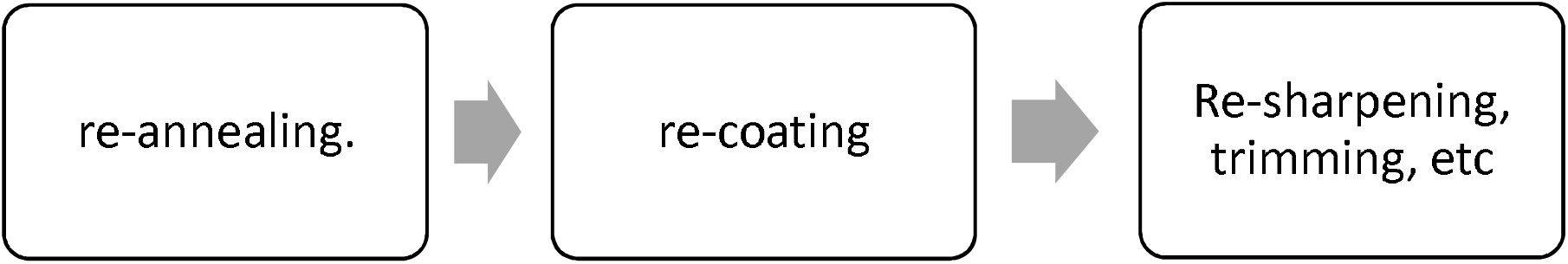
The proposed protocol for reconditioning neural microelectrodes with material failure

## Results and discussion

To demonstrate the effectiveness of the protocol proposed above, 27 cases of single-site neural microelectrodes were studied. The impedance of these tungsten electrodes had decreased, and they all had physical defects and needed to be removed. For this purpose, the electrodes that were bent at the tip were cut off. After washing them with distilled water, they were placed in the setup shown in picture 3. Care must be taken to ensure that the tip of the electrode, which we want to sharpen, is free of any contamination, including the remaining insulation on this surface. Using the holder, place the electrodes near the surface of the solution and in front of the camera. Next, manually adjust the starting point of the solution surface so that it is set near the resin on the upper part of the electrode (this is done using a camera and tester’s eye). From here on, activate automatic settings and first transfer the solution level to the solution holding chamber through the pump. After a set time, actuate the pump motor in the opposite direction and transfer the solution in the opposite direction to the main chamber. The solution level will rise with the tip of the electrode. The suggested time interval for adjusting the phase of changes in the solution level is 5 to 10 seconds in each phase.

Figure 4 shows three examples of sharpened electrodes. The first image from the left is the image after cutting the tip of the electrode. The bottom images are the electrodes after etching. To improve the impedance of these electrodes, it was necessary to cover the tips of the electrodes with insulation. It is important to note that this insulation must be biocompatible. Therefore, we used a method where the insulator is heated up to 400 degrees Celsius until it becomes a liquid. Then, the tip of the electrode is dipped in it and spread on its tip. Finally, using a heater that is placed towards the tip of the electrode, the insulation is directed from the tip to its back to create a smooth surface on the surface of the electrode. To check the result, you should examine the smoothness of the surface of the electrode tip using a microscope.

**Figure 3:**
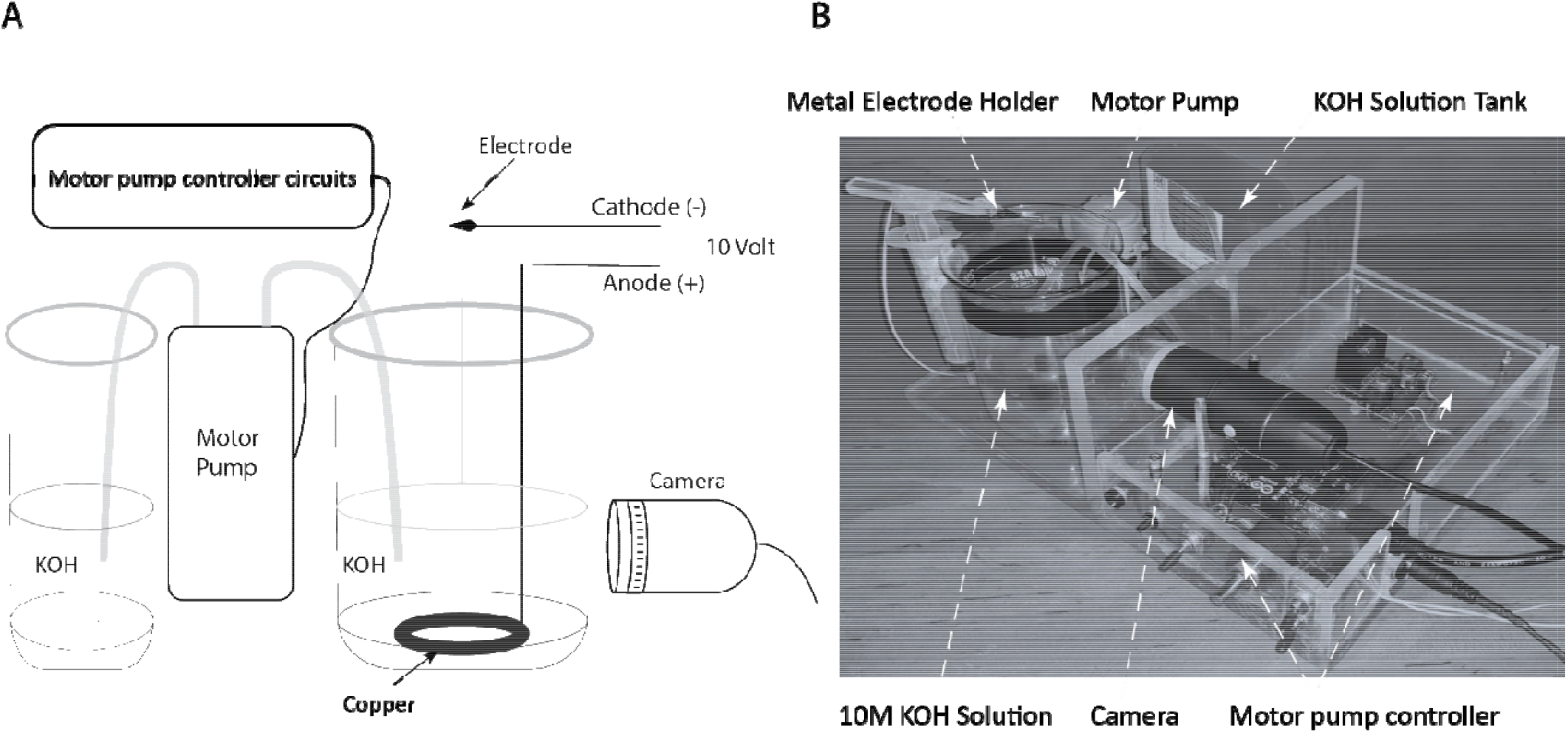
Electrode etching setup. A) Electrode etching setup schematic. In this setup, there is a tank of 10 M KOH solution in which a piece of copper metal is connected to a 10 V source with a communication wire. The other part of this circuit is the electrode which is fixed with a clamp. The part of the tip that we want to sharpen is placed in the solution and the end of the electrode is connected to the cathode of the DC source. A pump motor and another tank are used to change the level of the solution in the main tank by a few millimeters. Changes in the solution level are determined using controller circuits. Additionally, the changes in the level of the solution and the degree of sharpening of the electrode tip can be monitored through a camera. B) Real Image taken from the setup All parts of setup were determined in order.

**Figure 4:**
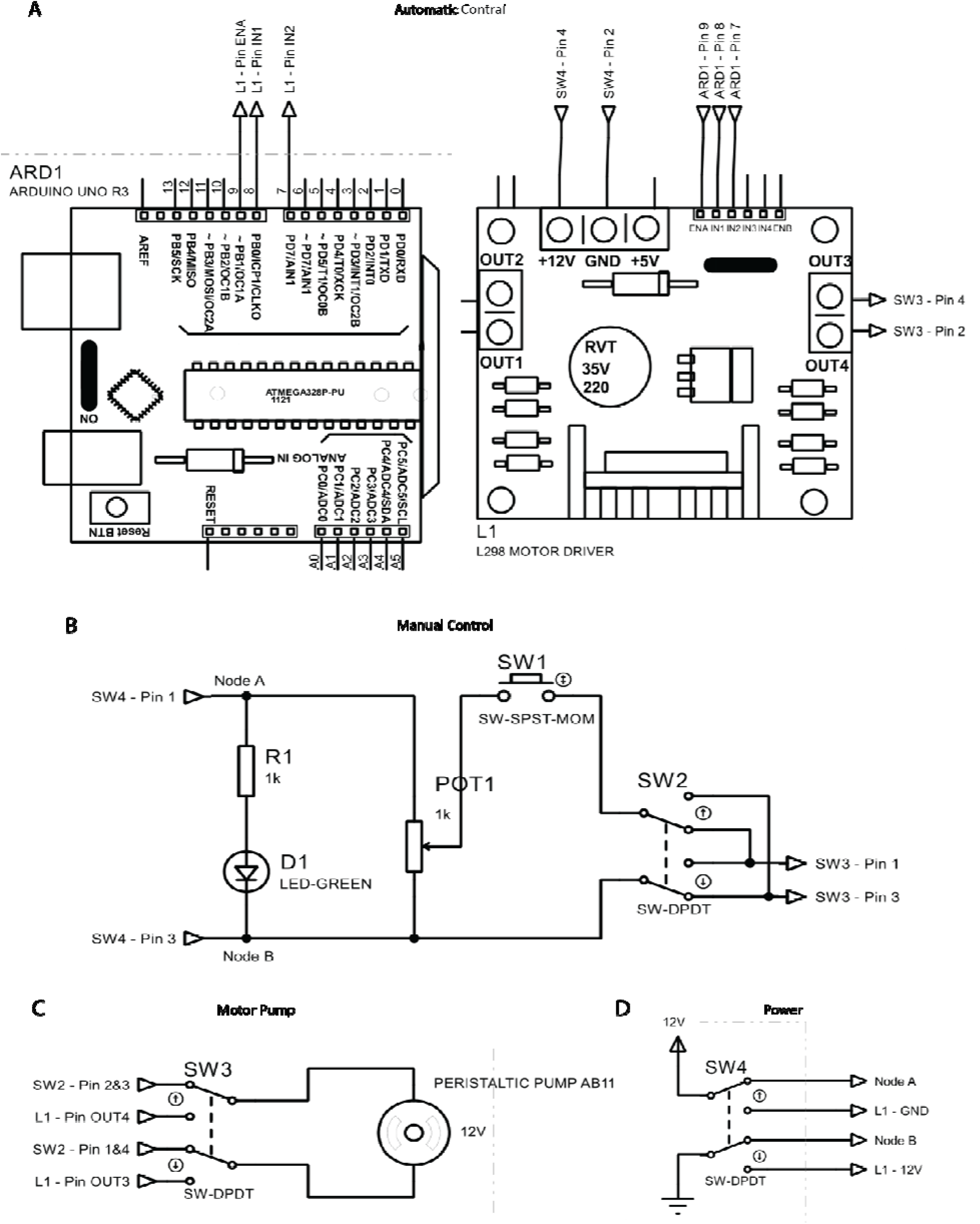
Motor pump controller circuits. A) The automatic control system consists of two main components: an Arduino UNO board that is used to program and set the automatic operation of the system, and a L298N driver module that is used to start the motor. Since the motor draws a lot of current which can damage the Arduino board chip, we need to use the driver module to start the motor. B) The manual control system includes four main components: a reset button that is used to turn on the motor, a 6-pin two-position DPDT switch that is used to determine the direction of motor rotation, a 1kΩ multi-train potentiometer that is used to adjust the motor speed, and an LED that is in series with a 1kΩ resistor and is used to indicate when manual control mode is enabled. C) The pump motor block consists of two main components: a 12-volt peristaltic liquid pump motor and a 6-pin two-mode DPDT switch that is used to determine whether the motor receives input from the automatic control block or the manual control block. D) The power supply block includes two main components: a 12-volt power input jack that is used to supply power to the setup and a 6-pin dual-mode DPDT switch that is used to determine the power supply to the automatic control block or the manual control block.

After measuring the impedance of all the tested electrodes, as seen in Figure 5, 26 of the electrodes had their impedance decreased compared to before applying our proposed protocol, and the impedances have been improved with noticeable results.

**Figure 5:**
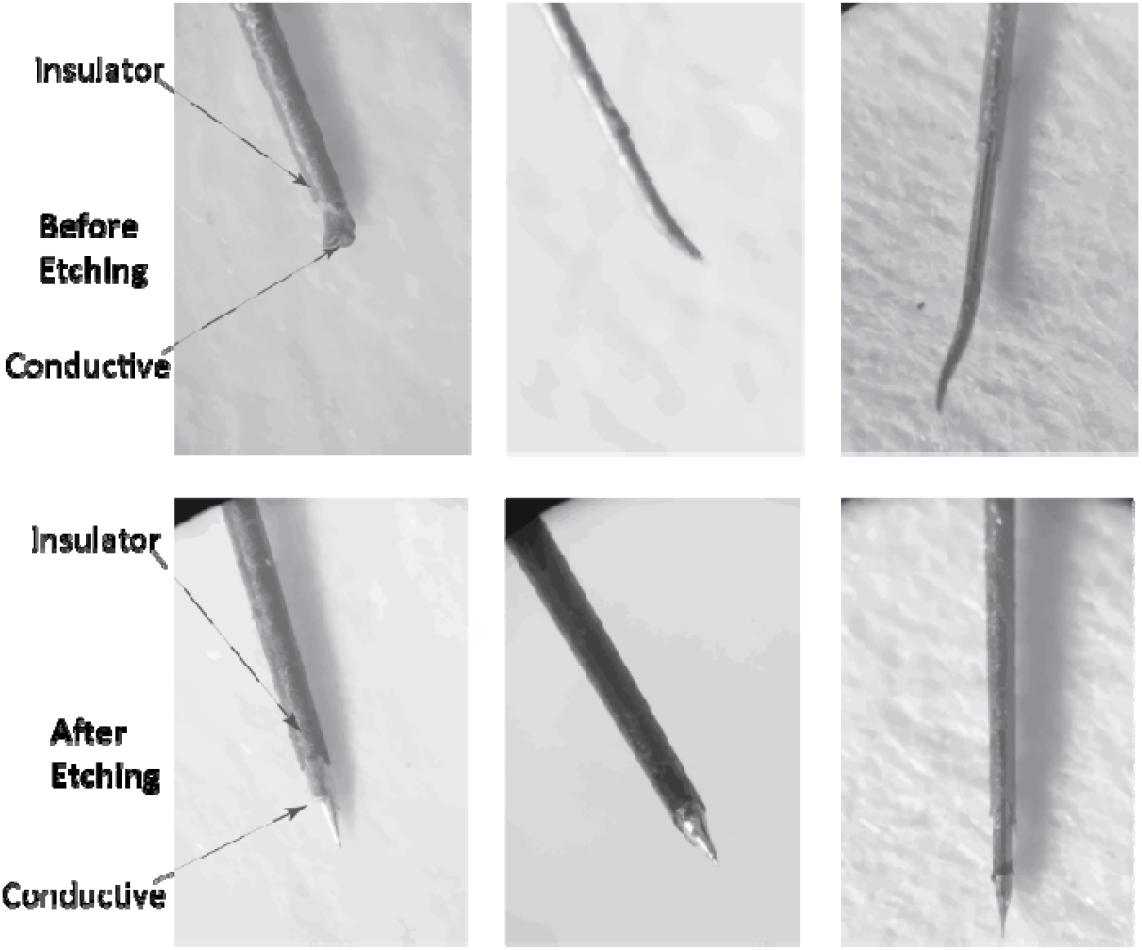
Microscopic images of the tips of three electrode samples before and after etching.

**Figure 6.**
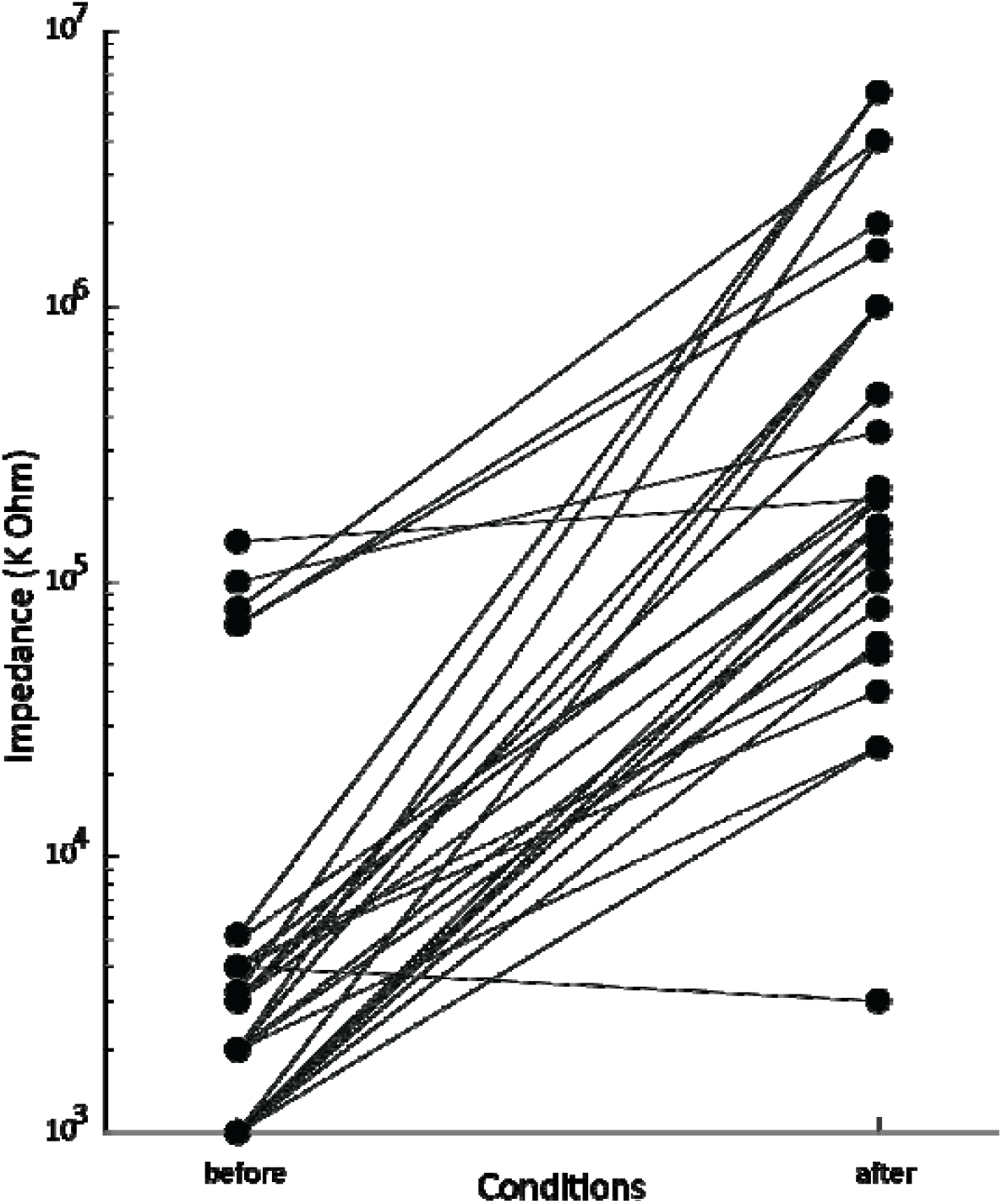
Impedance value of electrodes before and after etching. Most of electrodes impedance were recovered.

Figure 4 shows three examples of sharpened electrodes. The first image from the left is the image after cutting the tip of the electrode. The bottom images are the electrodes after etching. To improve the impedance of these electrodes, it was necessary to cover the tips of the electrodes with insulation. It is important to note that this insulation must be biocompatible. Therefore, we used a method where the insulator is heated up to 400 degrees Celsius until it becomes a liquid. Then, the tip of the electrode is dipped in it and spread on its tip. Finally, using a heater that is placed towards the tip of the electrode, the insulation is directed from the tip to its back to create a smooth surface on the surface of the electrode. To check the result, you should examine the smoothness of the surface of the electrode tip using a microscope.

After measuring the impedance of all the tested electrodes, as seen in Figure 5, 26 of the electrodes had their impedance decreased compared to before applying our proposed protocol, and the impedances have been improved with noticeable results.

## Conclusion

The study presented the effectiveness of a proposed protocol for reconditioning tungsten electrodes. The study involved 27 cases of single-site neural microelectrodes, and the results showed that the impedance of 26 of the electrodes had decreased compared to before applying the protocol. The protocol also describes the methods used to recondition the electrodes, including re-annealing and re-coating, and the process of sharpening the electrodes. The insulation used to cover the tips of the electrodes must be biocompatible, and the smoothness of the surface of the electrode tip should be examined using a microscope. The study concludes by stating that the proposed protocol was effective in improving the impedance of the electrodes. The methods used to recondition the electrodes and sharpen them were described in detail. The protocol provides valuable information for researchers and professionals working in the field of neural microelectrodes. Using simple and low-cost methods, this issue can be resolved, and costs can be saved.

